# Scalable and robust phylogenetic tree reconstruction from copy-number data with Sparse Rooted Neighbor Joining

**DOI:** 10.64898/2026.07.30.739152

**Authors:** Vittorio Zampinetti, Harald Melin, Alva Hallin, Jens Lagergren

## Abstract

**Background:** Phylogenetic tree reconstruction from single cell data based on copy-number alterations (CNAs) is an important problem in cancer genomics. Methods have been developed to address this problem by computing pairwise distances between copy-number profiles and employing a tree reconstruction algorithm. Despite the tight interplay between distance estimation and tree reconstruction, these two steps are often treated as separate problems, with the choice of the reconstruction algorithm receiving little attention. Most methods rely on classical Neighbor Joining (NJ), an algorithm designed for unrooted phylogenies that does not account for the fixed diploid root inherent to copy-number evolution.

**Results:** We identify the Deepest Least Common Ancestor NJ (DLCA–NJ), not previously applied in this context, as the appropriate algorithm for phylogenies from copy-number data. By leveraging the known diploid root, it consistently outperforms standard NJ on simulated benchmarks across all evaluated metrics, with the most pronounced improvement in root placement accuracy. Building on these findings, we introduce Sparse Rooted Neighbor Joining (SRNJ), a scalable adaptation of DLCA–NJ. SRNJ significantly reduces running time while trading off only a minor loss in accuracy. We provide theoretical and empirical evidence of robustness to mutation rate using both synthetic and real biological datasets.

**Conclusions:** Rooted NJ variants offer a principled way to exploit the known diploid root when reconstructing phylogenies from copy-number data, and SRNJ extends this advantage to datasets whose size places the full distance matrix out of reach. The gains are clearest where distances are reliable, as on simulated data, while on real data accuracy appears to be constrained by distance estimation rather than by the reconstruction algorithm, leaving room for improvement as callers advance.

## 1 Introduction

Copy-number alterations (CNAs) are a hallmark of cancer genomes and play a crucial role in tumor evolution and heterogeneity. Single-cell DNA sequencing technologies have enabled the profiling of CNAs at single-cell resolution, providing valuable insights nto the evolutionary history of tumors [1–4]. CNAs can be used to identify driver mutations, understand tumor progression, and inform treatment strategies [5, 6]. To this end, several computational methods have been developed to reconstruct phylogenetic trees from CNA data, addressing challenges such as noise, missing data, and complex events like whole-genome duplications (WGDs) [7–13].

Just like in classical phylogenetics, the most scalable methods for tree reconstruction from CNA data are distance-based, called so because they rely only on pairwise distances between samples to infer the topology, where a sample can be either a copy-number profile or a sequence of raw DNA read-counts. In particular, these methods are based on two key components, namely the estimation of pairwise distances between samples, and the application of a tree reconstruction algorithm that takes these distances as input and infers the tree topology by optimizing some criterion. For the sake of clarity, the latter will be hereafter referred to as *distance-based* method, whereas the combination of the two will be referred to as *sequence-based* method, following the naming convention of [14]. While often treated as sequential, the two steps of a sequence-based method are fundamentally linked and may overlap. Early distance-based methods use simple distance estimates such as the Euclidean or Manhattan distance between copy-number (CN) profiles [3, 4], which must be considered a poor approximation of the underlying evolutionary process, given that copy-number alterations affect contiguous genomic regions, introducing strong dependencies between adjacent bins. More advanced distances include the Minimal Event Distance (MED), implemented in MEDICC2 [10]. This widely used approach computes the minimum number of copy-number alteration events required to transform one copy-number profile into another and uses this quantity for phylogenetic reconstruction. As such, MED is based on a minimum-event criterion rather than an explicit probabilistic model of CNA evolution. In classical phylogenetics, model-based distances account for the possibility that multiple evolutionary events may affect the same genomic locus over time, providing a statistical framework for distance estimation under an assumed evolutionary process [15]. Such models can improve phylogenetic inference in settings where recurrent or overlapping mutational events are common [16, 17]. For classical substitution models these model-corrected distances are *additive*: the distance between any two taxa equals the sum of the edge lengths along the path joining them in the tree, the property that distance-based reconstruction relies on. An uncorrected raw distance may instead deviate from additivity and bias the resulting trees [16, 17]. Among copy-number methods, SCONCE2 [13] is, to our knowledge, the only one that attempts an analogous correction, implementing a probabilistic model of copy-number evolution and using it to estimate a triplet of distance measures between two sequences: *t*_1_ (distance from the root to the Least Common Ancestor, or LCA), and *t*_2_*, t*_3_ (distances from the LCA to the two leaves). Unlike the classical case, however, these model-based estimates are not guaranteed to be additive on the copy-number tree; we make this distinction precise in Appendix A.1. Despite this rich output, it is assumed that the tree reconstruction should proceed by building a distance matrix with *t*_2_ + *t*_3_ as the eaf-to-leaf distance and then applying a standard tree reconstruction algorithm, such as Neighbor Joining (NJ) [18]. This approach disregards *t*_1_, the root-to-LCA distance, which carries valuable information about the divergence of the two leaves from the known root and can therefore be used to improve tree reconstruction.

Standard NJ reconstructs unrooted trees from leaf-to-leaf distances and therefore t does not utilize the known ancestral root. This is a significant limitation in cancer genomics, as the healthy diploid state provides a natural root for the tumor phylogeny. While root information can be incorporated by treating the root as an outgroup, this may lead to errors when the outgroup is distant [19–21]. Specialized algorithms exist that incorporate root information directly: Asymmetric NJ (ANJ) [22] uses distances from leaves to the LCA, while Deepest Least Common Ancestor Neighbor Joining (DLCA–NJ) [23] utilizes root-to-LCA distances. We find that DLCA–NJ, although not previously used for cancer phylogenetics, is better suited to this context than standard NJ. It constructs the tree directly from the root using the root-to-LCA distance, which corresponds exactly to the *t*_1_ distance estimated by SCONCE2. This makes DLCA–NJ a natural choice for tree reconstruction from copy-number data.

As data availability increases in both sample size *n* and sequence length *m* (i.e., the number of genomic bins), the estimation of the full pairwise distance matrix becomes the primary computational bottleneck. Let *O*(*f* (*m*)) be the time complexity of computing the distance between a single pair of sequences. While simple distance estimators are linear in *m*, modern probabilistic models explicitly account for hidden evolutionary events to increase accuracy, at the cost of substantially higher per-pair complexity. For SCONCE2, for example, computing a single pairwise distance requires *O*(*mK*^4^) time, where *K* is the maximum copy-number state (typically set to 10), making the full *O*(*n*^2^*mK*^4^) matrix computation the dominant bottleneck regardless of the relative sizes of *m* and *n* [13, 24].

In the specific context of copy-number evolution, current methods primarily mitigate these computational costs by segmenting genomic bins through breakpoint inference [9, 10, 12]. This reduces the effective dimension of *m*, although the benefit may be limited when noisy or highly fragmented copy-number profiles lead to the inference of many spurious breakpoints, which inflate the effective dimension and propagate calling error into the distance estimates [14]. In contrast, Sparse Neighbor Joining (SNJ) [24] avoids the estimation of all *n*^2^ pairwise distances by identifying optimal merges while computing only a logarithmic fraction of the distance matrix, maintaining theoretical guarantees with a reduced complexity of *O*(*f* (*m*)*n* log *n*).

In this work, we demonstrate that replacing traditional NJ with DLCA–NJ significantly improves reconstruction accuracy for methods that use model-based distances such as those from SCONCE2, with the largest gains in root placement metrics. Furthermore, we adapt the SNJ approach to the rooted setting, deriving the *Sparse Rooted Neighbor Joining* (SRNJ). This algorithm extends the DLCA framework to large datasets by avoiding full matrix computation, achieving a worst-case time complexity of *O*(*f* (*m*)*n* log *n*). The remainder of this paper is organized as follows. In Section 2, we provide the theoretical grounds for rooted tree reconstruction and introduce the SRNJ method and its properties. In Section 3, we present benchmarking experiments on synthetic and real datasets to evaluate the accuracy and execution time of the proposed methods, followed by a discussion of the results and current limitations. Finally, we conclude with a summary of our findings and future research directions in Section 4.

## 2 Materials and Methods

### 2.1 Deepest-LCA Neighbor Joining

In classical phylogenetics, distance-based algorithms reconstruct a tree topology *T* from a non-negative distance matrix *D* defined over *n* observations referred to as *taxa* (singular: taxon). These taxa may represent any distinct biological units or data profiles, such as genomic sequences or read-count profiles. Recall from the introduction that a distance matrix is *additive* when each entry equals the sum of the edge lengths along the path connecting the two taxa in a tree. The most widely used tree reconstruction algorithm is Neighbor Joining [18], which iteratively merges pairs of taxa that minimize the *Q* criterion defined as

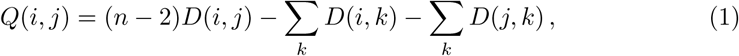

where *D*(*i, j*) is the distance between taxa *i* and *j*. If *D* is additive for a tree *T*, then NJ reconstructs *T* exactly. Since in practice most distance matrices are noisy estimates of the underlying additive matrix, research has characterized the robustness of NJ to errors in the distance matrix. In particular, it has been shown that NJ always reconstructs the true tree if the error in the distance matrix is less than half of the minimum edge length in the tree, or in other words, the *l*_∞_*-radius* of NJ is 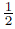. Another crucial result is that NJ correctly reconstructs all edges in the true tree *T* that are onger than 4 times the maximum error in the distance matrix; NJ is said to have *edge l_∞_-radius* of 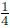 [25, 26].

While in classical phylogenetics the ancestral sequence at the root is typically unknown, cancer phylogenetics offers a distinct advantage. Because tumor cells evolve from a healthy diploid genome, the root of the tree is effectively known: it corresponds to the diploid state, i.e., a uniform copy number of 2 across all genomic loci. This property allows distance estimation methods to calculate the distance from the root to each observed taxon, providing an additional source of information beyond just the distances between taxa. Given a fixed root, the distance from the root to the LCA of two taxa provides an alternative measure to the leaf-to-leaf distance. Following the terminology in [23], we refer to this distance as the LCA distance. Let *L* be the matrix consisting of estimated root-to-LCA distances for each pair of taxa. Then one can show that if *D* is additive for a tree *T* with root *r*, then the matrix *L* defined as

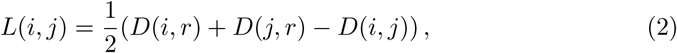

consists of the true distances from the root to the LCAs in *T*. However, LCA distances can also be estimated directly from triplets of sequences, using, e.g., maximum likelihood techniques, potentially leading to more accurate estimates [27, 28]. More generally, we can say that any non-negative matrix *L* over a set *S* is an LCA-matrix if *L*(*i, i*) = max*_j∈S_L*(*i, j*) for all *i* ∈ *S* and *L*(*i, j*) ≥ min(*L*(*i, k*)*, L*(*j, k*)) for all *i, j, k* ∈ *S* (three-point condition). The DLCA–NJ algorithm reconstructs a rooted tree from an LCA-matrix *L* by iteratively merging pairs of taxa with the maximum (deepest) LCA distance, hence its name. After a merge, the LCA-matrix is updated by means of a reduction formula which has to be defined in a way that preserves the additivity. For example, any reduction formula of the form

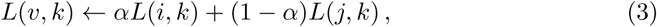

where *v* is the new node obtained by merging *i* and *j*, and with *α* ∈ [0, 1], maintains the additivity of the LCA-matrix. Notably, DLCA–NJ has been shown to be more robust than NJ in terms of edge *ℓ_∞_*-radius, with an optimal radius of 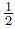 instead of the 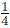 of NJ [23]. Although this property does not mean that DLCA–NJ is more accurate than NJ in general terms, it suggests that exploiting the known root can lead to improved reconstruction of short edges in phylogenies from copy-number data.

### 2.2 Sparse Rooted NJ

Standard DLCA–NJ, like NJ, requires the prior computation of the full *n* × *n* LCA distance matrix and runs in *O*(*n*^3^) time. However, the cost of computing the distance matrix typically dominates the overall running time of the sequence-based method. To mitigate this, we adapt the Sparse NJ framework [24] to the rooted case, deriving SRNJ.

SRNJ bypasses the full matrix computation by selecting taxa to compare based on the current state of the reconstructed tree. The algorithm starts by building a rooted backbone tree using DLCA–NJ on a subset of *k*_0_ taxa, small enough such that the full LCA distance matrix can be computed in *O*(*f* (*m*)*n* log *n*) time (e.g., 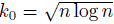). Then, it iteratively finds the edges where to insert the remaining leaves by exploring the tree and calculating the distances as needed. This strategy ensures that only *O*(*n* log *n*) distances are computed, while preserving the consistency of DLCA–NJ.

#### 2.2.1 Overview of SRNJ

Let *T^k^*^0^ be the rooted binary backbone tree built on a subset of *k*_0_ ≥ 2 leaves for which the full LCA distance matrix is computed. At each iteration *k* = *k*_0_ + 1*, …, n*, a new leaf *l_k_* is added to the current tree *T^k−^*^1^ by finding the correct edge for its insertion. The algorithm first identifies a *centroid* node in *T^k−^*^1^, chosen so that, if removed, the largest connected component has at most half of the *k* − 1 leaves. The existence of such a node is ensured by the Centroid Decomposition Theorem [29]. Then, two *orienting* leaves, *l_a_, l_b_*, are selected at random, one from each of the two subtrees below the centroid, i.e., following the edges pointing away from the root. The LCA distance between the new leaf *l_k_* and each of the two orienting leaves is computed, and the DLCA–NJ tree among these three leaves is built to determine which of the three components to explore next. Two cases can occur: either *l_k_* forms a cherry with one of the orienting leaves, or *l_a_, l_b_* form a cherry and *l_k_* is placed as sibling of their parent. In the first case, *l_k_* appears to be closer to one of the orienting leaves than to the other, and therefore the algorithm explores the component containing the orienting leaf that forms the cherry, while in the second case, it explores the component that contains the root. In both cases, the current centroid becomes a *barrier* node. A barrier node isolates the edges remaining to be explored. While the leaves beyond this node remain available as potential orienting leaves, the barrier effectively represents them during the next centroid selection step, ensuring that the algorithm explores the tree in a balanced way, i.e., it only counts as one leaf for the purpose of selecting the next centroid. Since at each step the algorithm recurses into a component with fewer than half the current leaves, the search depth is at most log_2_(*k* − 1) for a tree of *k* − 1 leaves, yielding the *O*(log *n*) distance computations per insertion. Whenever the recursion is called on a subtree that only contains one leaf, one barrier or the root node, the new leaf *l_k_*is inserted on the edge connecting such node to the current centroid. While the SRNJ algorithm focuses exclusively on recovering the tree topology, edge lengths can be estimated in a subsequent post-processing step using standard distance-based optimization criteria such as ordinary or weighted least squares [15]. A schematic representation of the algorithm is shown in Figure 1.

**Fig. 1.**
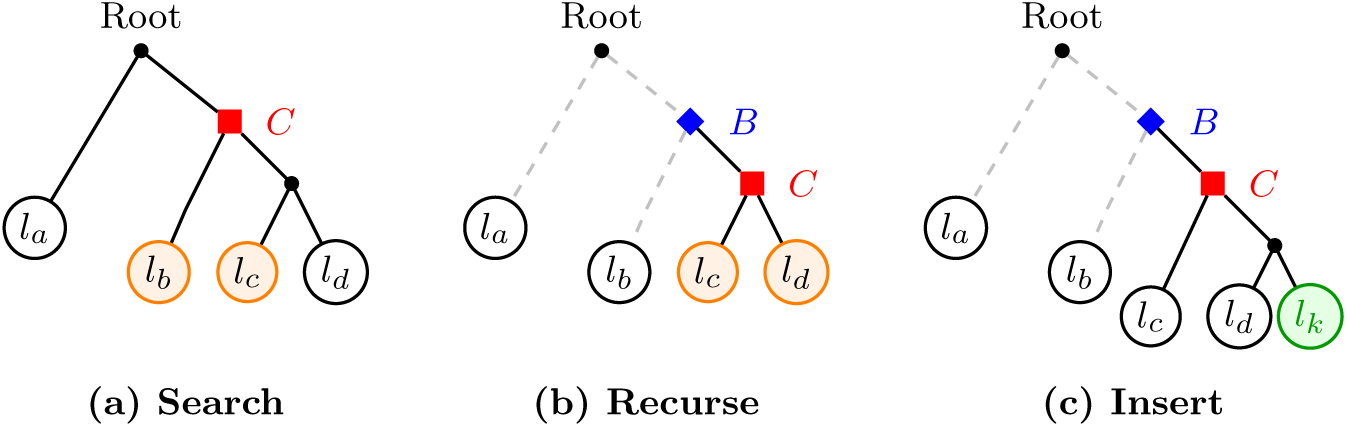
SRNJ iteration steps. (a) A centroid *C* is chosen and orienting leaves *l_b_, l_c_* are selected. (b) After comparing them with *l_k_*, the algorithm recurses into the *l_c_* subtree; the old centroid becomes a barrier *B* and a new *C* is chosen, with *l_c_, l_d_* as orienting leaves. (c) The recursion continues towards the *l_d_* subtree, at which point, with only one leaf left, *l_k_* is inserted on the edge connecting *C* to *l_d_*.

#### 2.2.2 Selection of orienting leaves

The selection of orienting leaves is a crucial step in the algorithm, as it determines the accuracy of the direction chosen at each recursion. When the distance *f* (*m*) is expensive to compute (e.g., *O*(*mK*^4^) for SCONCE2), we subsample log *n* candidates at random from each subtree and evaluate only those. Scoring log *n* candidates at each of the *O*(log *n*) recursion levels increases the total number of distance evaluations to *O*(*f* (*m*) *n* log^2^ *n*), still substantially lower than the *O*(*f* (*m*) *n*^2^) full-matrix cost.

Alternatively, when the pairwise distance can be computed in *O*(*m*) time with a small constant—for instance the Hamming distance on rounded copy-number profiles, or the NLL distance (see Appendix A.2)—all leaves in each subtree can be scored without materially increasing the overall cost, and no subsampling is needed.

For both regimes, the orienting leaf can be selected either as the candidate with the smallest leaf-to-leaf distance or, following the logic of DLCA–NJ, as the one with the argest estimated LCA depth. We evaluate these variants empirically in Section 3.4. Crucially, this heuristic changes only *which* candidates are scored, not the insertion rule applied afterwards: on an additive matrix any valid pair of orienting leaves yields the correct local topology, so subsampling preserves the consistency guarantee of Section 2.3.1 and affects only the constant and logarithmic factors in the running time, leaving the *O*(*f* (*m*) *n* log *n*) bound of the original sparse method unchanged. The cheap-proxy regime is likewise a scoring device only; the NLL distance it can use s defined in Appendix A.2.

### 2.3 Theoretical properties of SRNJ

#### 2.3.1 Consistency

Here we use *consistency* in the sense of the reconstruction algorithm: a method is consistent if it recovers the true tree exactly whenever the input distance matrix is additive, regardless of the distance estimator used to produce it.

DLCA–NJ is consistent: if *D* = *D_T_* is additive for a tree *T*, the algorithm reconstructs *T* exactly. SRNJ invokes DLCA–NJ only on selected triplets of *D*. If *D* is additive, every submatrix is also additive, and DLCA–NJ returns the correct local topology at each insertion step. Hence each recursive subtree selection follows the true path toward the correct edge. By induction over the insertion order, every taxon is inserted at the correct edge, and SRNJ reconstructs *T* exactly. Therefore SRNJ is consistent.

#### 2.3.2 Robustness

The greedy insertion strategy of SRNJ means it does not inherit the edge *ℓ_∞_*-radius guarantee of DLCA–NJ. At each insertion step only a local quartet is evaluated; errors on short edges can therefore compound across iterations in ways that a global algorithm avoids by averaging over the full distance matrix. As a consequence, short edges are more susceptible to misplacement by SRNJ than by DLCA–NJ. This limitation can be mitigated by considering multiple orienting leaves at each iteration, as described above, increasing the likelihood of selecting the correct subtree for exploration, and is empirically evaluated in Section 3.4.

#### 2.3.3 Complexity Analysis

The time complexity of the algorithm is *O*(*f* (*m*)*n* log *n*), where *f* (*m*) is the time complexity of computing the distance between two sequences of length *m*. In fact, the algorithm computes the full distance matrix for a subset of 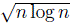 leaves to build the backbone tree, which takes *O*(*f* (*m*)*n* log *n*) time. Then, for each of the remaining 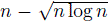 leaves, it computes the distance to at most 2 log *n* new leaves, which also takes *O*(*f* (*m*)*n* log *n*) time in total. While the iterative nature of the algorithm limits global parallelization compared to traditional Neighbor-Joining, this is more than compensated for by the reduction in total distance calculations from *O*(*n*^2^) to *O*(*n* log *n*). Parallelization can still be effectively applied to the backbone matrix construction and the independent distance evaluations required for each leaf insertion.

### 2.4 Evaluation metrics

To evaluate the accuracy of the reconstructed trees, we use four commonly used metrics in phylogenetics: the Robinson-Foulds (RF) distance [30], the quartet distance [31], the transfer distance [32] and the triplet distance [33], which is specifically designed for rooted trees. The RF distance measures the number of splits (bipartitions of the leaf set) that are present in one tree but not in the other. In particular, we use the normalized RF distance, which is the RF distance divided by the maximum possible RF distance for trees with the given number of leaves i.e.,

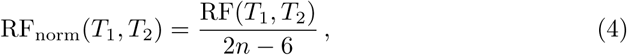

where *n* is the number of leaves in the trees. This allows us to compare the accuracy of the reconstructed trees across different datasets with varying number of leaves. The RF distance is widely used due to its simplicity and computational efficiency. However, t only measures whether splits are identical or not, and therefore does not capture how different two incompatible splits are. As a consequence, small topological changes (e.g., moving a single taxon) can alter several splits and lead to a relatively large RF distance, while large rearrangements that affect the same set of splits are penalized equally. Moreover, because the number of splits grows linearly with the number of eaves, the RF distance often takes values close to its maximum for moderately large trees, reducing its ability to discriminate between different levels of topological error. The quartet distance, on the other hand, counts the number of quartets (sets of four eaves) whose induced topology differs between the two trees, reported as a fraction of all 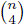 quartets. Because this fraction is taken over a number of quartets that grows much faster than the number of splits underlying the RF distance, it varies more smoothly and provides a more fine-grained comparison between trees with a large number of leaves [34]. The transfer distance [32] provides a more fine-grained comparison than the RF distance by measuring how similar individual edges are between two trees. For an edge *b^∗^* in the reference tree *T_ref_*, the transfer index is defined as the minimum Hamming distance between the bipartition induced by *b^∗^* and any bipartition induced by an edge *b* in the estimated tree *T_est_*. Intuitively, this quantity corresponds to the minimum number of leaves that must be moved from one side of the bipartition to the other to make an edge in *T_est_* match *b^∗^*. To obtain a single score for the whole tree, we compute the transfer index for each internal edge of *T_ref_* and report their arithmetic mean. Additionally, as a way of assessing the accuracy of root placement, we compute the transfer index only for the edge connecting the root to its children in *T_ref_* and normalize it by the maximum number of leaves that can be transferred, which is the minimum number of leaves on either side of the bipartition minus one. We denote this metric the *root-split* distance. The triplet distance [33] (introduced as *triples* distance in the original paper) is the rooted analog of the quartet distance. It considers all *^n^* subsets of three leaves and for each subset {*i, j, k*}, the metric determines if the rooted topological relationship is consistent between the reference and the estimated tree (e.g., (*i,* (*j, k*)) ̸= ((*i, j*)*, k*)). The triplet distance is defined as the number of such subsets with differing topologies, normalized by the total number of possible triplets. Because the hierarchical nesting of a triple is determined by the path to the root, this metric is highly sensitive to root misplacement, so—together with the root-split distance defined above—it serves as an indicator of root-placement accuracy. All distances except the transfer distance are always normalized to take values between 0 and 1, even if not specifically mentioned in the text. Tree distances were computed using standard implementations: Robinson-Foulds via *DendroPy* [35], quartet and triplet distances via *tqDist* [36], and transfer distance via *Booster* [37].

## 3 Results and Discussion

### 3.1 Benchmark on *in-house* simulated data

We benchmarked the algorithms on data generated with a simple *in-house* CN evoution model, which allows us to compute the LCA distances using the ground truth simulated CN sequences, including those of ancestral sequences, thereby avoiding CN calling errors. Trees are generated with a birth-death process (birth–rate=1) and the number of CNAs on each edge is sampled from a Poisson distribution with mean *λ* = 5. Each CNA event represents a gain or a loss (therefore changing the copy number by 1) of a segment whose start position is drawn uniformly over the bins and whose ength is sampled from another Poisson with mean *µ* = 200 bins. The number of bins s fixed to *m* = 1000 and the number of leaves varies in *N* ∈ {20, 50, 100, 200, 500}. An example of such simulated data is shown in Figure A1 in the Appendix. We run NJ, DLCA–NJ, SNJ and SRNJ on the same dataset and compare the reconstructed trees with the true tree using the metrics described above. The distances are computed from the ancestral CN profiles by means of a simple evolutionary model similar to the one introduced in [11] (see Appendix A.1 for details); for each pair of leaves, we compute three distances (root-to-LCA and the two LCA-to-leaf distances) from the known CN profiles. Using the known ancestral profiles makes this an idealized setting that isolates the reconstruction algorithm from distance-estimation error and thus provides an upper bound on achievable accuracy; the effect of estimating distances from data instead is examined in the subsequent experiments (Section 3.2). We note that the distance computation, while accounting for dependencies in the copy-number profiles, is a simplified evolutionary model (with the Markov property) and does not perfectly match the generative process used for simulation, which is biologically more realistic. The distances are therefore not guaranteed to be additive on the true tree, and as such, some residual reconstruction error is expected even when ancestral profiles are known exactly. We evaluate the Sparse Rooted NJ algorithm using three variants that differ in the size of the orienting-leaf candidate pool, selecting in all cases the candidate with the smallest leaf-to-leaf distance. SRNJ-1 uses a single random orienting leaf, and SRNJ-log samples log *n* random candidates. SRNJ-all scores its candidates—all inserted leaves—with a cheap surrogate distance rather than with the expensive estimator (e.g., SCONCE2) used by SRNJ-log. Its full surrogate distance matrix is precomputed once, before SRNJ runs, at a cost of *αmN* ^2^, which we assume to be far smaller than SRNJ’s running time. These variants allow us to assess the impact of the candidate-pool size on the accuracy of the reconstructed trees, and to evaluate the trade-off between accuracy and computational cost. A broader comparison of orienting leaf selection strategies, including proxy-distance approaches, is presented in Section 3.4. For comparison, we also execute the original SNJ algorithm using the smallest leaf-to-leaf distance selection criterion on log *n* random candidates, as described in the original paper. Additionally, we include ANJ [22] and FastME [38] in the comparison for a more comprehensive benchmark. ANJ is included as another rooted NJ variant that uses leaf-to-LCA distances rather than root-to-LCA distances, allowing us to isolate the benefit of the specific LCA formulation used by DLCA–NJ. FastME combines NJ with heuristic tree search strategies to optimize the balanced minimum evolution criterion, without any guarantee of consistency, but with the same time complexity of NJ, making it a valid alternative to NJ. The results in Figure 2 show that DLCA–NJ outperforms NJ in the quartet, triplet and transfer distances across all dataset sizes, with the improvement being more pronounced for larger datasets. In terms of RF distance, DLCA–NJ is better than or comparable to NJ, with the gap closing for the largest datasets. FastME shows the best accuracy in terms of RF distance, but it is outperformed by DLCA–NJ in terms of quartet and triplet distance, which indicate a better placement of the root for the latter. This effect is even more evident in the root-split distance (i.e., accuracy of the root bipartition) shown in Figure 3, where only DLCA–NJ and the SRNJ variants reach perfect accuracy, while NJ, SNJ, ANJ and FastME misplace the root increasingly often as the number of leaves grows. SRNJ shows a significant improvement over its NJ counterpart (SNJ) with respect to all metrics, with a pronounced improvement in RF and transfer distance when more than a single orienting-leaf candidate is used (SRNJ-log and SRNJ-all). Compared to DLCA–NJ, with larger datasets, SRNJ shows particularly lower performance in terms of RF distance. This might be due to a combination of the greedy insertion strategy and the limited resolution of the RF distance, as the other metrics do not show such a pronounced drop in performance. Notably, the performance of SRNJ in terms of quartet and triplet distance is inferior only to that of DLCA–NJ, despite the greedy nsertion. This suggests that SRNJ effectively recovers local nested relationships and sub-clade structures better than the other algorithms not based on LCA distances, even if some global topological errors are introduced by the greedy insertion strategy.

**Fig. 2.**
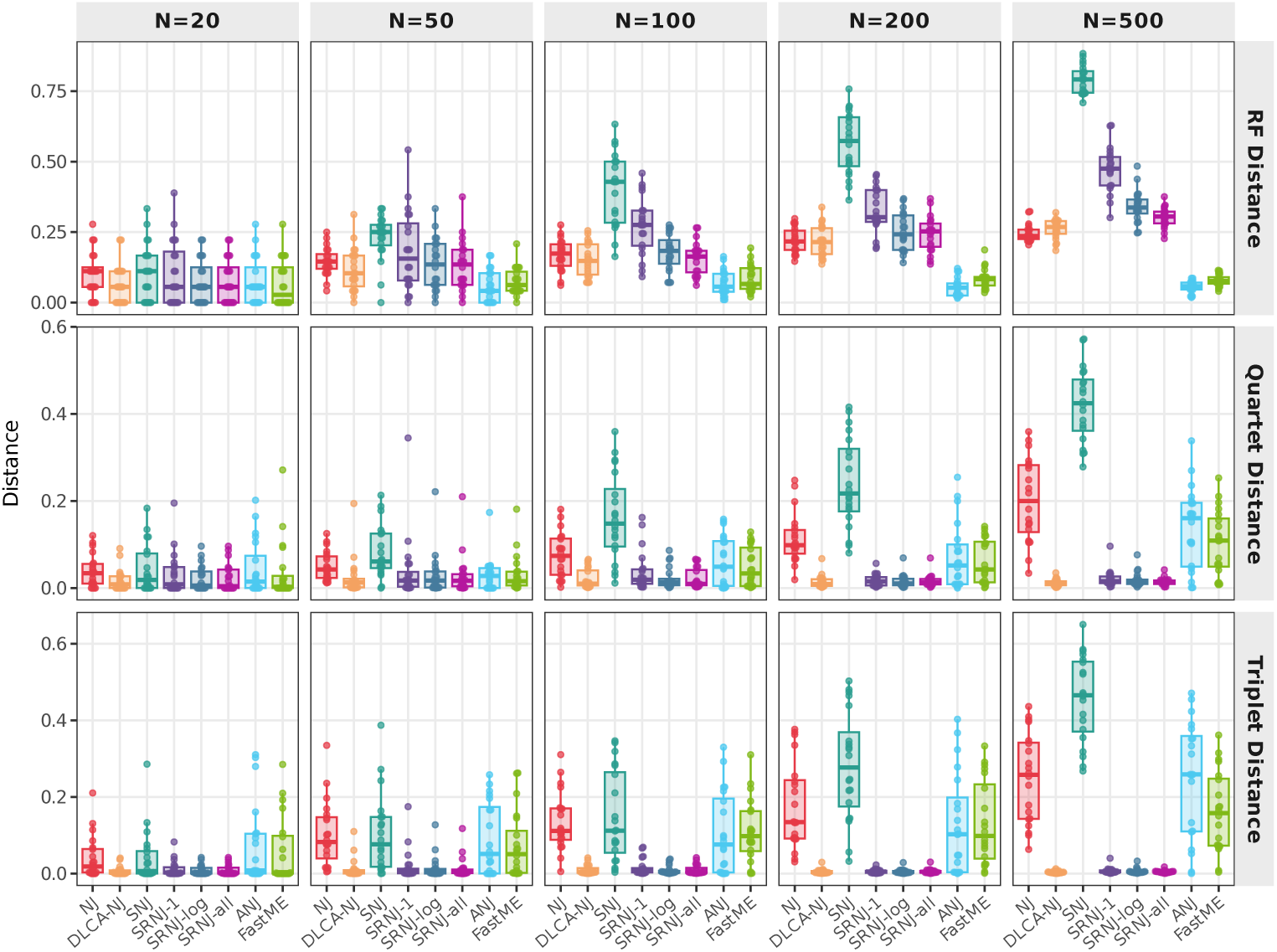
Reconstruction accuracy on *in-house* simulated data. Boxplot aggregate scores over 20 different data seeds, for *N ∈ {*20, 50, 100, 200, 500*}* leaves (columns) and the RF, quartet and triplet distances (rows; lower is better). SRNJ is executed in three variants that all select the orienting leaf with the smallest leaf-to-leaf distance but differ in the candidate-pool size: a single random leaf (SRNJ-1), log *n* random candidates (SRNJ-log), or all inserted leaves with NLL proxy (SRNJ-all).

**Fig. 3.**
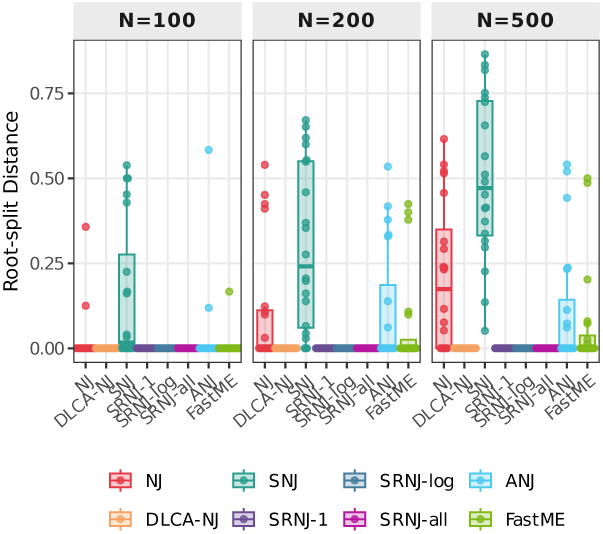
Root placement accuracy on *in-house*simulated data. Boxplots showing the *root-split* distance in the experiment with *in-house* simulated data. Distances are computed from ground-truth ancestral copy-number profiles using the JC-breakpoint correction (Appendix A.1).

### 3.2 Realistic CNA simulation and distance estimation

For a more realistic benchmark where CN sequences are not known, we evaluated the performance of DLCA–NJ and SRNJ in combination with distances estimated via the state-of-the-art methods SCONCE2 and MEDICC2. We simulated datasets of different sizes, with *N* ∈ {50, 100} leaves, and fixed sequence length of *m* = 1000 genomic bins, equally divided into 10 chromosomes. A dataset consists of a tree topology and a set of sequences at the leaves, where each sequence contains the read-counts for each genomic bin and is generated according to the CN profile of the corresponding eaf. The simulated datasets were generated with CNAsim [39], a simulator of copy-number evolution that allows us to generate trees with varying numbers of leaves and mutation rates. We set the average number of CNAs per edge to *λ* ∈ {2, 3}, which is a realistic scenario for cancer evolution, especially in the case of punctuated evolution, where a single cell can carry from many tens to hundreds of CNAs [3, 40].

We repeated the simulation for 20 replicates for each dataset size and ran SCONCE2 (with maximum copy-number state *K* = 10) on each dataset to estimate the *t*_1_, *t*_2_ and *t*_3_ distances, which were then used as input for NJ and DLCA–NJ. Like n the original SCONCE2 paper, we summed the *t*_2_ and *t*_3_ edge lengths to form a standard leaf-to-leaf distance matrix for NJ, while we adopted the *t*_1_ edge lengths as the root-to-LCA distances for DLCA–NJ. We include MEDICC2 in the comparison to assess the impact of using a model-based distance measure instead of the MED distance, which is not derived from an explicit probabilistic model of evolution. Since MEDICC2 requires copy-number profiles as input, we used the total copy numbers nferred by SCONCE2 for a fair comparison, and reconstructed the tree with NJ with outgroup rooting, as in the original MEDICC2 paper. We report the distribution of the quartet distance, RF distance, transfer distance and root-split distance between the true tree and the reconstructed trees for the different combinations of distance estimation methods and tree reconstruction algorithms in Figure 4(b) (see Figure A2 n the Appendix for a complete overview). Also, we report the per-replicate difference Δ = SCONCE2 − MEDICC2 for each reconstruction method in Figure 4(a); negative values indicate better performance with SCONCE2 distances.

**Fig. 4.**
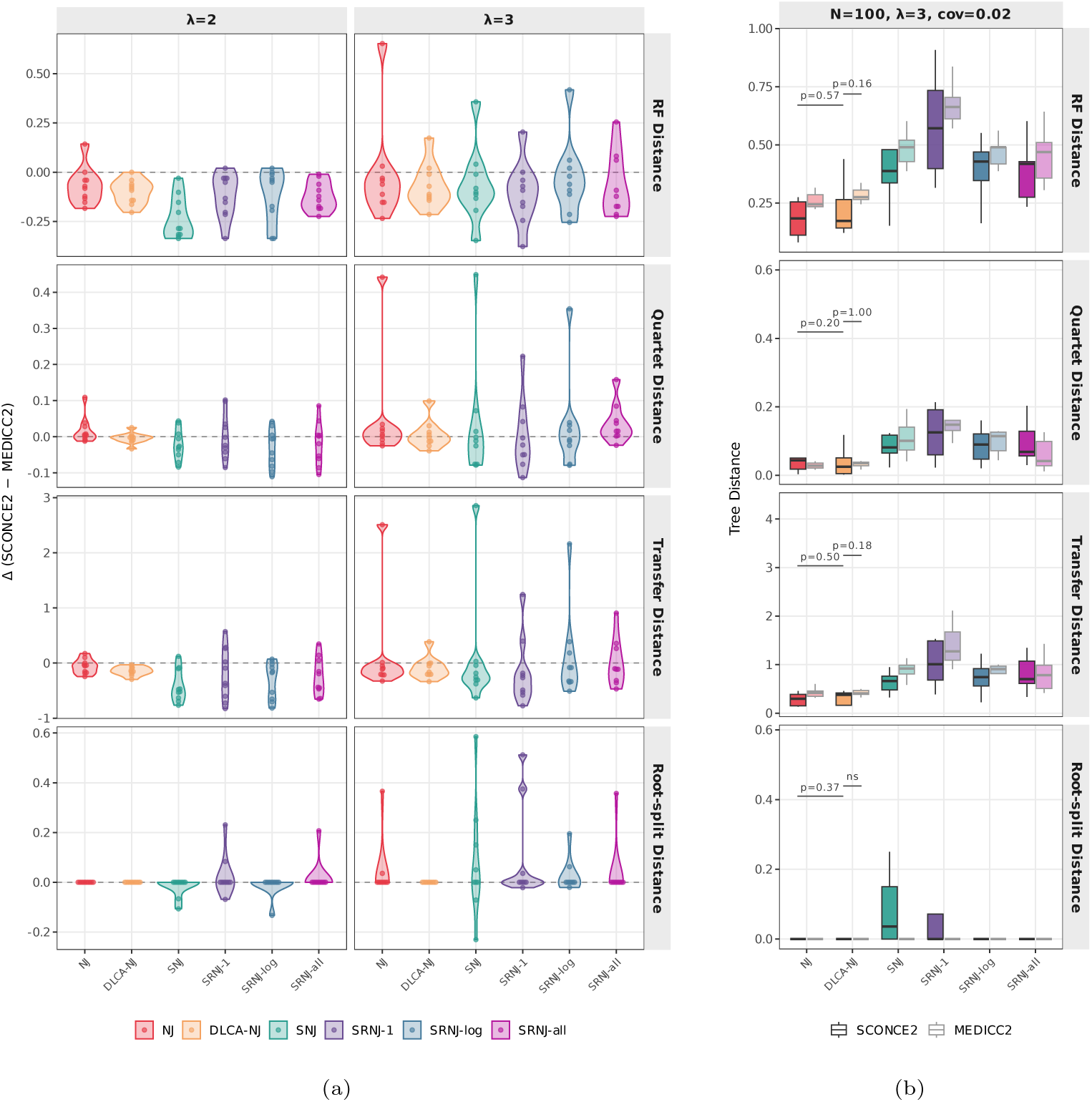
Accuracy with SCONCE2 and MEDICC2 distances on CNAsim data. Aggregated results of the experiment with synthetic data generated with CNAsim (20 replicates) and distances estimated with SCONCE2 and MEDICC2. Panel (a) shows the per-replicate difference Δ = SCONCE2 *−* MEDICC2 for each tree reconstruction method (NJ, DLCA–NJ, SNJ, SRNJ-1, SRNJ-log, SRNJ-all) on simulated datasets of *N* = 100 cells, across mutation rates *λ ∈ {*2, 3*}*. Rows show (top to bottom) RF distance, quartet distance, transfer distance and root-split distance between the true tree and the reconstructed tree. Negative values indicate better performance by SCONCE2. Panel (b) shows the tree accuracy for *λ* = 3, with SCONCE2 (opaque) and MEDICC2 (transparent) compared side by side. Brackets indicate paired Wilcoxon tests p-values. The three SRNJ variants all select the orienting leaf with the smallest leaf-to-leaf distance and differ in the candidate pool: SRNJ-1 (single random leaf), SRNJ-log (log *n* random candidates) and SRNJ-all (all inserted leaves).

We find that SCONCE2 distance estimates lead to more accurate trees than MEDICC2 distances: paired with DLCA–NJ, SCONCE2 distances significantly reduce the reconstruction error over MEDICC2 distances. On these inferred distances, how-ever, the margin of DLCA–NJ over the plain NJ ordering is small and not statistically significant; the algorithmic benefit of root-aware ordering is isolated on exact distances, in the in-house benchmark and in the ground-truth control (Appendix A.5). In absolute terms, DLCA–NJ with SCONCE2 distances achieves the best performance, ndicating the need for both a model-based distance estimation and a tree reconstruction algorithm that can leverage the root information. We also ran the three versions of Sparse Rooted NJ described in the previous experiment. We find that SRNJ achieves its best performance with the larger candidate pools (SRNJ-log and SRNJ-all, which are close to each other) and, although slightly less accurate than the dense DLCA– NJ, it inherits the same benefit of the model-based SCONCE2 distances over MED distances and improves on its own sparse baseline SNJ when considering the root-split distance; the single-candidate SRNJ-1 is consistently the weakest variant. On these inferred distances the SRNJ-over-SNJ margin is modest — the full advantage is recovered once copy-number-calling error is removed, as shown by the ground-truth control in Appendix A.5. Due to the long execution time of SCONCE2 (see Section 3.3), we were not able to run the experiment with larger datasets, but we expect the benefits of DLCA–NJ and SRNJ to be even more pronounced for larger trees, as shown in the previous experiment with *in-house* simulated data.

### 3.3 SRNJ running time

SRNJ is a sequence-based method that takes sequences as input and relies on the distance estimation procedure as a subroutine to compute distances for individual pairs only when needed. However, neither MEDICC2 nor SCONCE2 software implementations support pairwise distance queries. For the previous experiments we bypassed this issue by computing the full distance matrix and then only using the required distances for SRNJ, but this approach does not allow to evaluate the actual running time of SRNJ. Therefore, to compare the scalability of the different algorithms, we emulate the total running time by tracking the number of required distance evaluations when executing SNJ and SRNJ, and we multiply it by the empirical time required by SCONCE2 to compute a single distance. For this purpose, we timed the execution of SCONCE2 in the previous experiment for each dataset with *N* = 100 and estimated the mean time required per pairwise distance computation as the total time divided by the number of pairs (i.e., *n*(*n* − 1)*/*2), obtaining 220.5 ± 66.5 seconds (mean ± SD over 20 replicates). This approach allows for a comparison of the running times shown in Figure 5, directly highlighting how the reduced number of distance computations in SRNJ translates to significant gains in total core-hours.

**Fig. 5.**
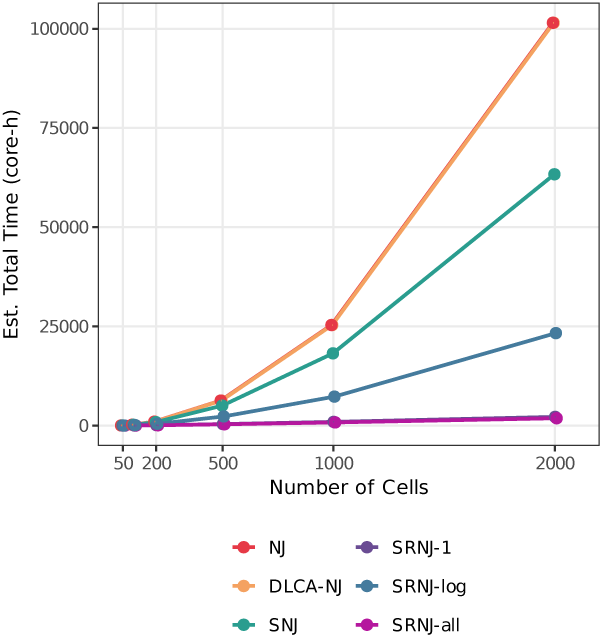
Estimated running time in core-hours. Estimated running time in core-hours for the different algorithms on simulated datasets of increasing size (*N* up to 2000 cells), obtained by multiplying the number of expensive (SCONCE2) pairwise distance evaluations by the measured per-pair cost. NJ and DLCA–NJ require the full *O*(*N* ^2^) distance matrix; SNJ and the SRNJ variants query far fewer pairs. SRNJ-all scores all candidate orienting leaves with a precomputed cheap distance matrix (computed once, *O*(*N* ^2^) but with a small constant and not requiring SCONCE2), so its number of expensive distance evaluations matches that of SRNJ-1 rather than growing with the candidate-pool size.

### 3.4 Effect of orienting leaf selection strategy

We evaluate how the choice of orienting leaf selection affects the accuracy of SRNJ across tree sizes *N* ∈ {20, 50, 100, 200, 500} using CNAsim-generated data (20 seeds per condition, *λ* = 5). Beyond the two built-in SRNJ criteria (min *d* and max LCA), we consider three additional families of selection strategies. The *ground-truth* (GT) oracle selects orienting leaves using the exact LCA distances computed from known ancestral sequences, providing an upper bound on the accuracy achievable by improved selection. The *Hamming* and *NLL* proxies are cheap surrogates derived from observed copy-number profiles and a healthy-cell baseline, requiring no additional tree inference steps.

Figure 6 shows that the choice of orienting leaf selection strategy has a pronounced effect on RF distance, especially for larger trees, while quartet and triplet distances are less sensitive to this choice. Among the SRNJ default strategies, the min-*d* criterion consistently outperforms max-LCA across all tree sizes and metrics. The GT oracle demonstrates that improving the selection of orienting leaves beyond the default SRNJ rules can substantially reduce the RF distance, approaching the accuracy of full DLCA–NJ at larger *N*. Notably, the Hamming and NLL proxy strategies achieve accuracy comparable to the GT oracle for RF distance, and both consistently outperform the default SRNJ sampling strategies (SRNJ-1, SRNJ-log, SRNJ-all), suggesting that scoring all candidate leaves with a numerically efficient distance is more effective than sampling a logarithmic subset with the exact distance. For quartet and triplet distances, all SRNJ strategies perform similarly, and the differences between selection criteria are small relative to the gap between SRNJ and DLCA–NJ. These results indicate that better orienting-leaf selection primarily reduces global topological errors (captured by RF distance), while local branching relationships are well recovered regardless of the strategy.

**Fig. 6.**
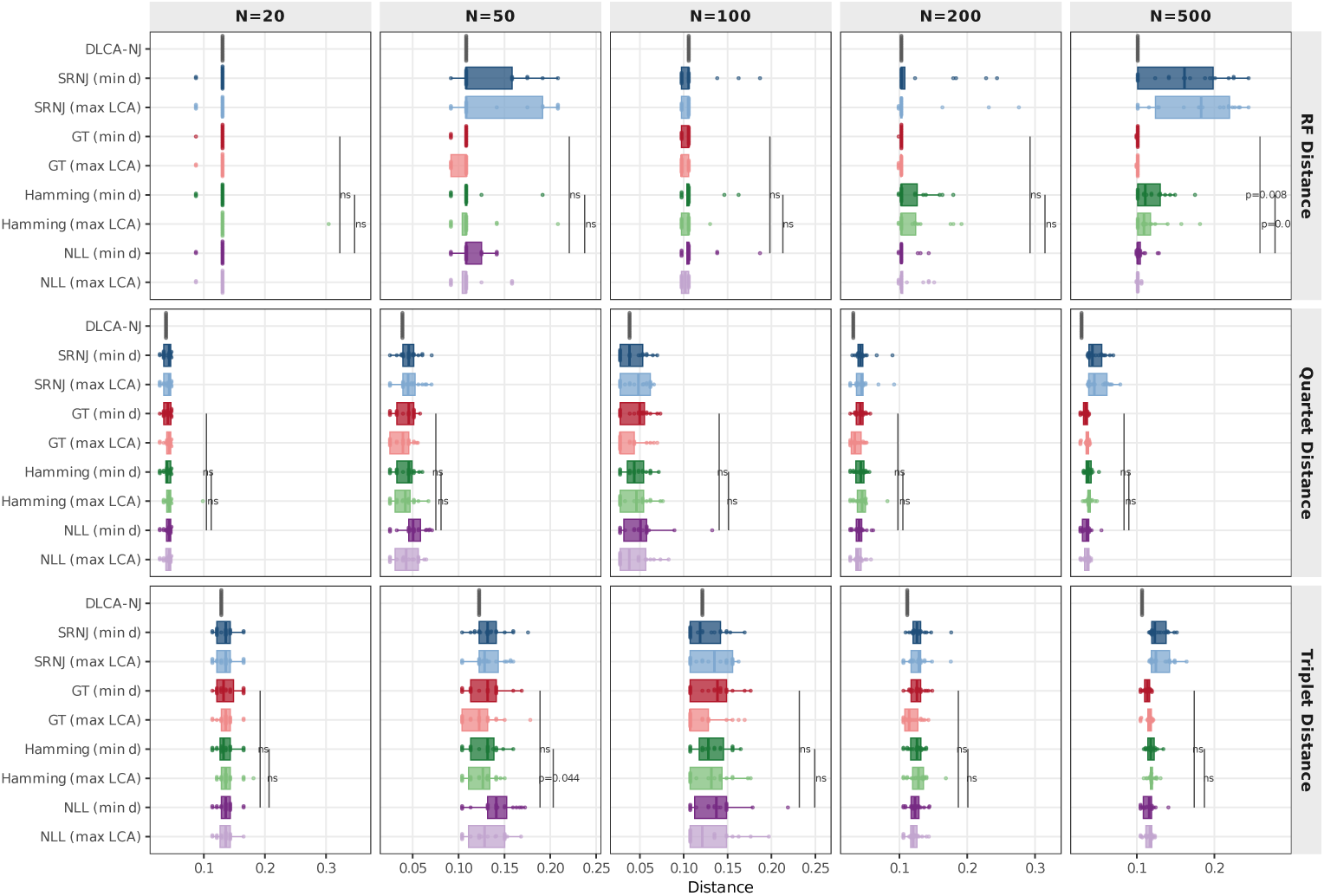
Orienting-leaf selection strategies on CNAsim data. Horizontal box plots (with individual seed points) of three tree-distance metrics (RF, quartet, and triplet distance) for nine orienting-eaf selection strategies applied to SRNJ, faceted by number of cells (*N ∈ {*20, 50, 100, 200, 500*}*, 20 seeds each). Strategies are grouped by distance source: baseline DLCA–NJ (grey), SRNJ defaults (blue), ground-truth oracle (red), Hamming proxy (green), and NLL proxy (purple). Within each group, two selection criteria are compared: *min-d* selects the orienting leaf with the smallest pair-wise cell-to-cell distance, while *max-LCA* follows the DLCA–NJ logic and selects the leaf whose LCA with the new taxon is estimated to be deepest (i.e., most distant from the root), using the available distance as a proxy. Lower tree distance indicates better reconstruction.

We note that on these CNAsim datasets the pairwise SNJ baseline performs comparably to the SRNJ variants, in contrast to the clear separation observed on the in-house data. As we show in Appendix A.5, this reflects the limited reliability of the SCONCE2-inferred distances rather than a genuine parity of the methods: on the very same CNAsim datasets, evaluated with ground-truth distances, the root-aware SRNJ again reconstructs the trees far more accurately than SNJ. The in-house benchmark, where distances are exact, is therefore the setting that isolates the algorithmic contribution of root information.

### 3.5 Breast cancer dataset

To evaluate tree reconstruction accuracy on real biological data, we apply DLCA–NJ, SRNJ and NJ to a breast cancer single-cell DNA dataset [41]. We compare the results with the clonal analysis reported in [42]. The dataset includes 10,000 cells from four tumor regions and one normal region. Due to the prohibitively high computational cost of running SCONCE2 on the full set, we focus on a subset of 500 cells from tumor region E, with bins of size 5Mb (*m* = 620). These cells were sampled so as to match the proportions of the eight clones (J–I to J–VIII) identified in the original CHISEL analysis. We use the normal region as diploid pseudo-bulk required for SCONCE2 distance estimation, and we set the maximum ploidy to *K* = 8. As a control, we compute a MEDICC2 distance matrix using the allele-specific copy-number profiles from CHISEL.

We evaluate how well the tree clades represent the clones inferred by CHISEL using the *clade-F1* score. For each clone, this metric identifies the clade with the highest F1 score and then computes the average of these scores across all clones. These clones are not a known ground truth but the output of the original CHISEL analysis, so the clade-F1 score should be read as agreement with a reference partition rather than with a true clonal structure. We run 50 bootstrap samples of 200 cells each from our 500-cell subset to quantify the sampling variability of this agreement across methods; this controls for variability due to cell subsampling but not for any systematic error in the reference clustering itself, which remains a limitation of the real-data evaluation. As shown in Figure 7, the allele-specific MEDICC2 distances yield substantially higher clade-F1 scores than SCONCE2 for every reconstruction method, a pattern consistent with reconstruction accuracy on this dataset being constrained by the inferred distances—which combine copy-number calling and branch-length estimation—rather than by the tree-building algorithm; separating these two contributions would require further experiments beyond the scope of this work. Within each distance source the reconstruction methods are close. With SCONCE2 distances, DLCA–NJ shows a small mprovement over NJ that does not reach statistical significance (paired Wilcoxon *p >* 0.1), and with MEDICC2 the two are essentially indistinguishable, reflecting the previous simulation experiment where the benefit of DLCA–NJ was more pronounced with model-based distances. The SRNJ algorithm falls slightly below the dense NJ and DLCA–NJ; the small F1 gap (of the order of 0.05) is significant but consistent with trading a small amount of accuracy for significant gains in scalability (Figure 7). We note that the clade-F1 score is insensitive to rooting and fine topology, the aspects where SRNJ’s root-aware reconstruction is most advantageous on ground truth, so this real-data metric does not reward exactly those strengths.

**Fig. 7.**
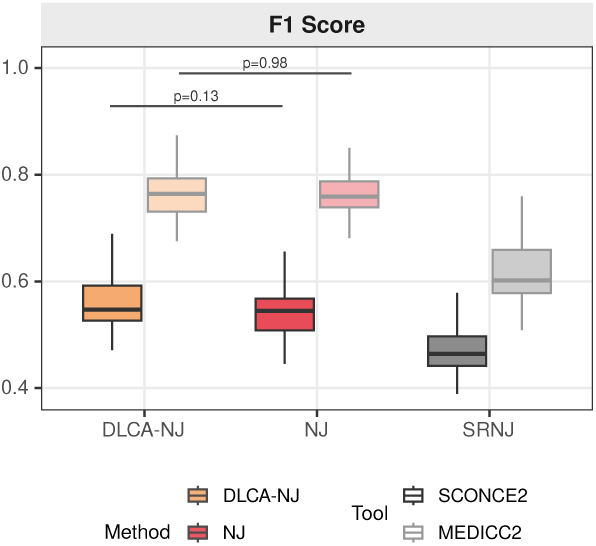
Clade-F1 score on the 10x breast cancer dataset. Boxplots showing the distribution of the clade-F1 score across 50 different bootstrap samples of 200 cells from region E of the 10x Genomics breast cancer dataset. The clade-F1 score is computed by comparing the clades in the reconstructed tree with the clones inferred in the original CHISEL analysis. Brackets indicate paired Wilcoxon test p-values.

### 3.6 Discussion

Taken together, these results highlight a gap between classical phylogenetics and the current copy-number evolution literature regarding the use of model-based distances. While corrected distances are standard in traditional phylogenetics, they remain uncommon in CNA studies. Our experiments suggest that model-based distance estimates, such as those provided by SCONCE2, can improve reconstruction accuracy, particularly under higher mutation rates and when combined with root-aware algorithms. Intuitively, a probabilistic model can account for hidden or overlapping events that a raw event count misses—the same rationale that motivates classical distance corrections—so these estimates move in the direction of a proper correction, even though in the copy-number setting their additivity is not established. The behaviour of SRNJ with accurate distances, as shown in the in-house benchmark, further suggests that rooted, iterative approaches preserve local branching orders and nested relationships more effectively than unrooted global methods.

A consideration that runs through these experiments is our reliance on SCONCE2 for copy-number calling. We are effectively constrained to it because it is, to our knowledge, the only method that natively outputs the decomposed root-to-LCA (*t*_1_) and eaf-to-LCA (*t*_2_*, t*_3_) distances that DLCA–NJ and SRNJ require; most callers provide only the copy-number profiles, from which these ancestral distances would still have to be estimated. The algorithms themselves are agnostic to the caller and would work with any future method that supplies, or permits the estimation of, these distances. We have therefore established the advantage of root-aware reconstruction on ground-truth and controlled simulations, where the distances are known or reliably estimated, and we make no claim of such an advantage in the real-data regime, where the qualty of copy-number inference dominates. Whether more accurate callers would expose the same root-aware advantage seen in simulation is an important open question that our experiments cannot settle.

## 4 Conclusion

In this work, we have demonstrated that incorporating root information through LCA distances improves phylogenetic tree reconstruction for CNA data. While the DLCA– NJ framework itself is not a contribution of this work, we established its specific suitability for CNA analysis, where the root is fixed and known. Building on it, we derived a novel algorithm, Sparse Rooted Neighbor Joining, which reduces the time complexity to *O*(*f* (*m*)*n* log *n*), where *f* (*m*) is the cost of a single distance estimation. We proved the consistency of SRNJ and evaluated its accuracy and robustness on both synthetic and real single-cell copy-number data.

The main limitation of SRNJ is practical rather than theoretical: existing distance estimation methods for copy-number evolution do not expose pairwise queries—they compute a full distance matrix at once rather than returning the distance of a single pair on demand, which is the access pattern SRNJ needs to realise its complexity advantage. Making these estimators more modular is therefore a practical prerequisite. A second limitation is that, on real data, reconstruction accuracy appears to be constrained by distance estimation (copy-number calling together with branch-length estimation) rather than by the reconstruction algorithm.

These limitations point to the natural directions for future work: adapting model-based distance estimation methods to answer individual pairwise queries, and improving the selection of orienting leaves through more advanced heuristics. More broadly, we hope this work encourages the development of probabilistic, model-based distance estimators such as SCONCE2. Their cost per pair has been a deterrent to their adoption, but that cost is multiplied by a quadratic number of pairs only for dense algorithms; under SRNJ the number of required estimates grows as *O*(*n* log *n*), so an estimator that is expensive but accurate becomes affordable at scales where computing a full distance matrix would not be. We believe this shifts the trade-off in favour of investing in better distance models, and provides guidance for future approaches to copy-number evolution and tumor phylogenetics.

## Declarations

### Ethics approval and consent to participate

Not applicable.

### Consent for publication

Not applicable.

### Availability of data and materials

The dataset supporting the conclusions of this article is publicly available from 10x Genomics [41], at https://cf.10xgenomics.com/samples/cell-dna/1.1.0/breast_tissue_aggr_10k/breast_tissue_aggr_10k_web_summary.html. All code required to reproduce the experimental results reported here is available in the SRNJ repository, https://github.com/Lagergren-Lab/srnj.

- Project name: SRNJ (Sparse Rooted Neighbor Joining)
- Project home page: https://github.com/Lagergren-Lab/srnj
- Archived version: https://doi.org/10.5281/zenodo.21398124
- Operating system(s): Platform independent
- Programming language: Python
- Other requirements: see the repository README for dependency versions
- License: GNU General Public License v3.0 (29 June 2007)
- Any restrictions to use by non-academics: None

### Authors’ contributions

VZ developed the method, implemented the software, designed and ran the experiments, analysed the results and wrote the manuscript. HM took part in project discussions and contributed ideas to the method. AH designed and ran the experiments comparing the orienting-leaf selection strategies. JL conceptualised and supervised the study. All authors read and approved the final manuscript.

## Appendix A Supplementary Material

### A.1 Simulation of *in-house* data

The *in-house* datasets used throughout the benchmark are generated by a single segment-based model of copy-number evolution. A cell lineage tree is drawn from a birth–death process (birth-rate 1), and each edge receives a number of CNA events sampled from a Poisson distribution with mean *λ* = 5, independently of the branch length. Each event is a gain or loss (changing the copy number by ±1) over a contiguous segment whose length is sampled from a Poisson with mean *µ* = 200 bins, starting from a diploid root over *m* = 1000 bins. Because every edge receives a comparable number of events, internal (shared) edges accumulate substantial ancestral signal, which is exactly what the root-aware methods exploit. Figures 2, 3 and A3 all use this same generative model.

We favour this in-house model over CNAsim for two reasons. First, it exposes the exact ancestral copy-number profiles at every internal node, so the root-to-LCA distances can be computed from the ground truth without any copy-number-calling error, isolating the behaviour of the tree-reconstruction algorithms. Second, it lets us control directly how copy-number evolution is distributed between internal (shared) and terminal (private) edges — the knob varied in the event-sharing experiment (Appendix A.4) — which CNAsim does not expose. CNAsim is instead used for the realistic distance-estimation experiments, where copy numbers are called from simulated reads.

**Fig. A1.**
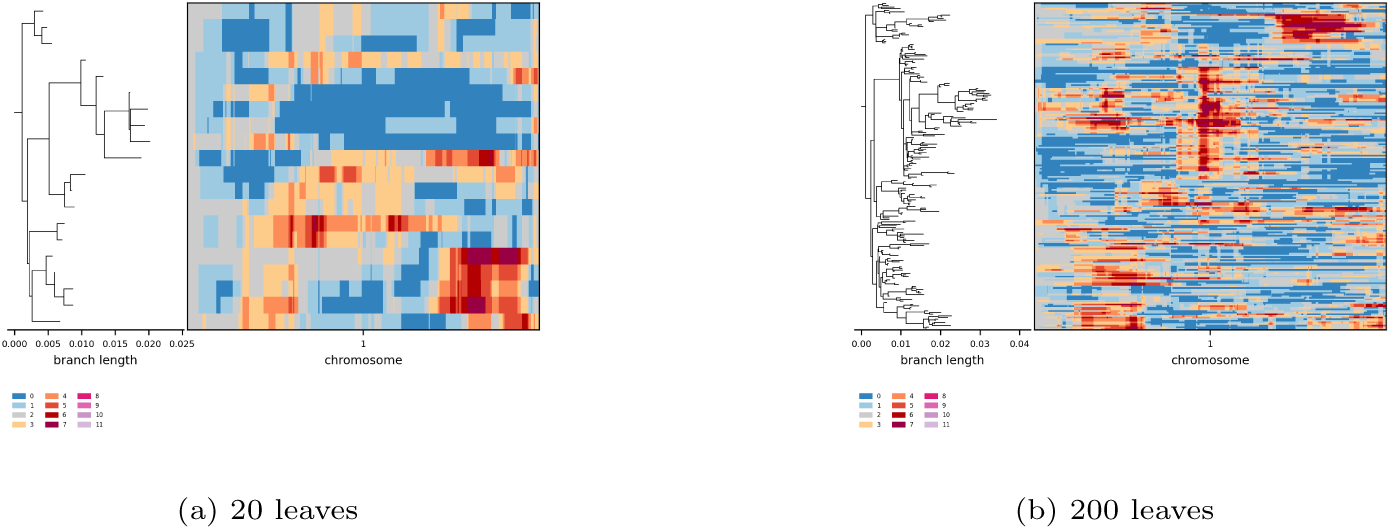
Example *in-house* simulated datasets. Example of two simulated datasets with the *in-house* model, featuring 20 (a) and 200 (b) leaves respectively.

#### Evolutionary model for distance correction

We first fix terminology, following classical phylogenetics. A *raw* distance is computed directly from the observed profiles. A *model-corrected* distance replaces that raw dissimilarity by an estimate obtained under a probabilistic model: given observations *x* and *y* and a family of distributions {*P_d_*} indexed by a distance parameter *d*, it is the maximum-likelihood estimate *d*^^^(*x, y*) = arg max*_d_ P_d_*(*x, y*). An *additive* distance is one that equals, for every pair, the sum of the edge lengths along the connecting path in some tree (the tree-metric property). Classical corrections, such as the Jukes–Cantor correction, are model-corrected *and* known to be additive. The distances used in this work—both the SCONCE2 estimates and the model below—are model-corrected in this statistical sense; we do not claim that they are additive on the true copy-number tree, and treat additivity as an open question.

To compute the corrected distances for the experiment with *in-house* simulated data, we use a simple evolutionary model of copy-number evolution similar to the one used in [11]. Specifically, we compute the mutation rate *p*^ over each edge in the tree as the number of breakpoints divided by the total number of bins. A breakpoint between two nodes *u, v* in the tree is detected over a pair of adjacent bins (*i, i* + 1) if

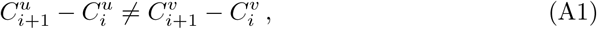

where 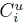 is the known copy number of node *u* at bin *i*. Then, we correct the distance taking into account the maximum allowed copy number *K* as follows:

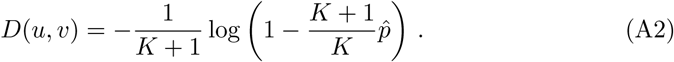

This correction is analogous to the Jukes-Cantor correction and easily derived by modeling the copy-number evolution as a continuous-time Markov process with *K* + 1 states (from 0 to *K*) and equal transition rates between any pair of states.

### A.2 Orienting-leaf selection

#### NLL distance

The NLL distance is a pairwise distance derived from how much each cell’s copy-number profile deviates from a healthy-cell baseline. Let **x**_1_, …, **x***_n_* be the normalised copy-number profiles of the tumor cells and **x***_n_*_+1_, …, **x***_n_*_+*m*0_ those of *m*_0_ normal cells. A per-bin Gaussian baseline is estimated as

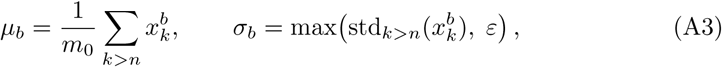

with *ε* = 10*^−^*^3^. Each cell *i* is scored at each bin *b* by the negative log-likelihood (NLL) under N (*µ_b_,* 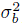):

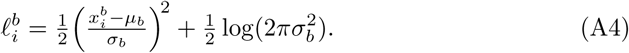

The pairwise NLL distance is then the mean absolute difference of these profiles:

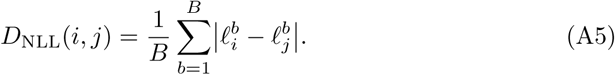

For the max-LCA variant, a root-distance proxy 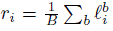 is used together with the additive-tree LCA decomposition *C_ij_* = (*r_i_* + *r_j_* − *D*_NLL_(*i, j*))*/*2 to estimate LCA depths. All operations are *O*(*m*) per pair with a small constant, making this distance suitable for scoring all candidate orienting leaves without a subsampling step.

### A.3 Additional results on CNAsim data

Figure A2 reports the complete results of the CNAsim experiment of Section 3, of which Figure 4 shows a representative subset: it spans all five tree distances, the three dataset sizes (*N* ∈ {20, 50, 100}) and both mutation rates (*λ* ∈ {2, 3}), for every combination of distance estimator and reconstruction algorithm. The trends discussed n the main text hold across these conditions. Trees reconstructed from SCONCE2 distances are generally more accurate than those obtained from MED distances; the dense algorithms (NJ and DLCA–NJ) retain a margin over their sparse counterparts; and among the SRNJ variants the single-candidate SRNJ-1 is consistently the weakest, while SRNJ-log and SRNJ-all remain close to one another. Reconstruction error grows with the number of cells for all methods. Notably, the root-split distance is close to zero for nearly every method and condition, so on these inferred distances it does not separate the rooted from the unrooted algorithms; the rooting advantage is instead apparent where the distances are exact (Figures 3 and A3).

### A.4 Effect of event sharing on reconstruction accuracy

The root-to-LCA formulation is expected to help most when copy-number evolution s dominated by early, shared events, so that internal (shared) edges carry strong ancestral signal, and to help least when evolution is mostly private to individual ineages. To probe this dependence directly, we vary the amount of copy-number evolution placed on internal (shared) edges of the *in-house* simulated trees using the same segment model as the main benchmark (Appendix A.1): terminal edges keep the base rate *λ* = 5, while internal edges receive a Poisson number of events with a controlled mean *λ*_int_ that we sweep from 0 (star-like trees, evolution almost entirely private) upward (*N* = 100 cells, 20 replicates). Figure A3 reports the reconstruction accuracy as a function of the realised mean number of shared events per internal edge. When shared ancestral signal is scarce, all methods struggle to recover the internal structure; as the number of shared events grows, the LCA-based methods (DLCA– NJ and SRNJ) increasingly outperform NJ and, especially, SNJ, confirming that the root-to-LCA formulation is most beneficial in the shared-evolution regime typical of punctuated tumor clonal expansions. The separation is clearest in the triplet and root-split distances (the rooting-sensitive metrics), where DLCA–NJ and SRNJ approach zero error while SNJ remains high; SRNJ also stays well below SNJ in RF distance across the whole range.

**Fig. A2.**
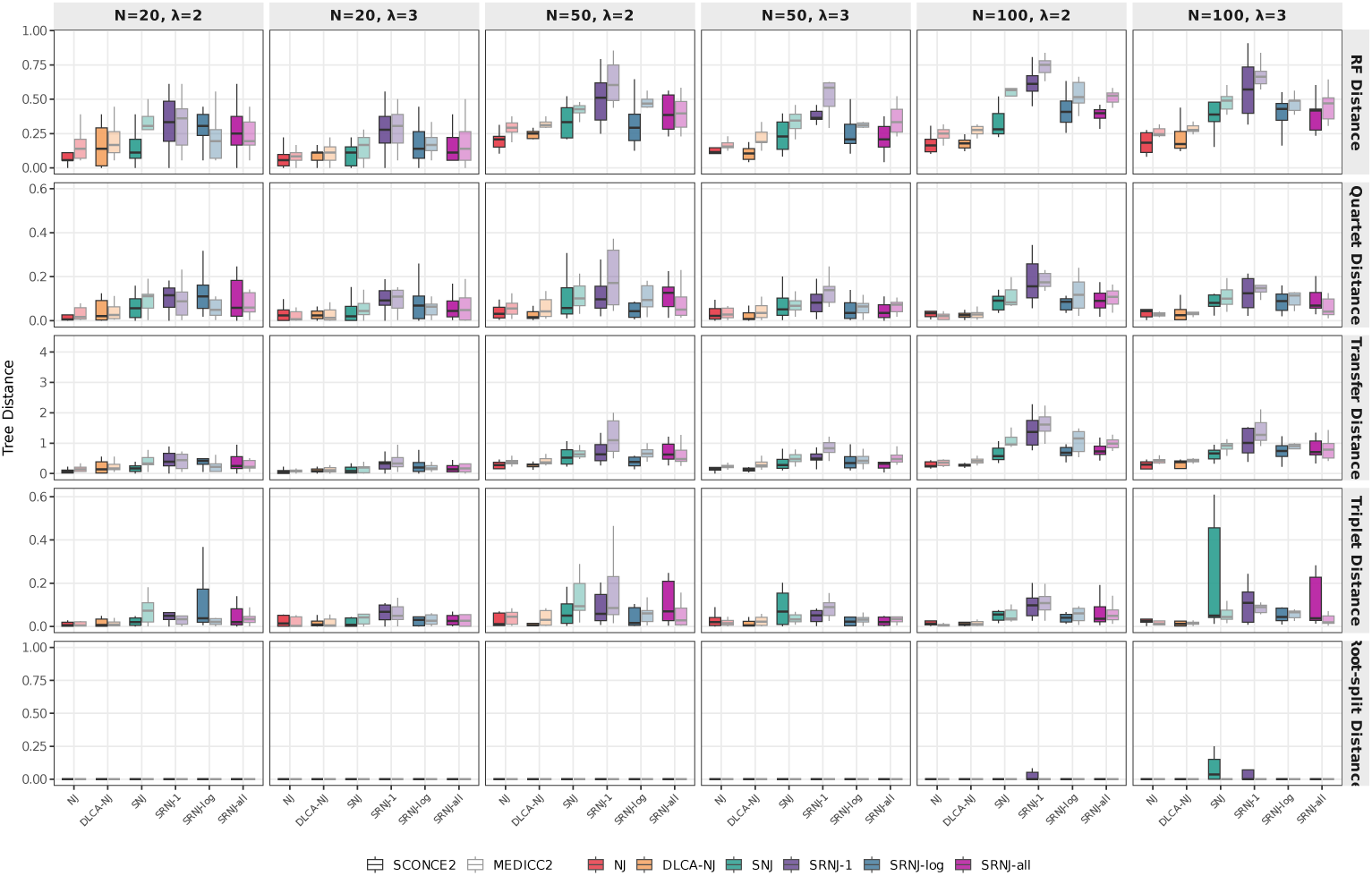
Complete tree-distance results on CNAsim data. Boxplots of tree distances between the true tree and reconstructed trees on **CNAsim-simulated data** across all evaluated conditions. Rows show (top to bottom) RF, quartet, transfer, triplet and root-split distance. Columns correspond to combinations of dataset size (*N ∈ {*20, 50, 100*}* cells) and mutation rate (*λ ∈ {*2, 3*}*). Each panel compares six tree reconstruction methods (NJ, DLCA–NJ, SNJ, SRNJ-1, SRNJ-log, SRNJ-all) applied to copy-number distances computed by SCONCE2 (opaque) and MEDICC2 (transparent). The SRNJ variants differ in the orienting-leaf candidate pool (single random leaf, log *n* random candidates, all inserted leaves). The sequencing coverage is 0.02*×*.

**Fig. A3.**
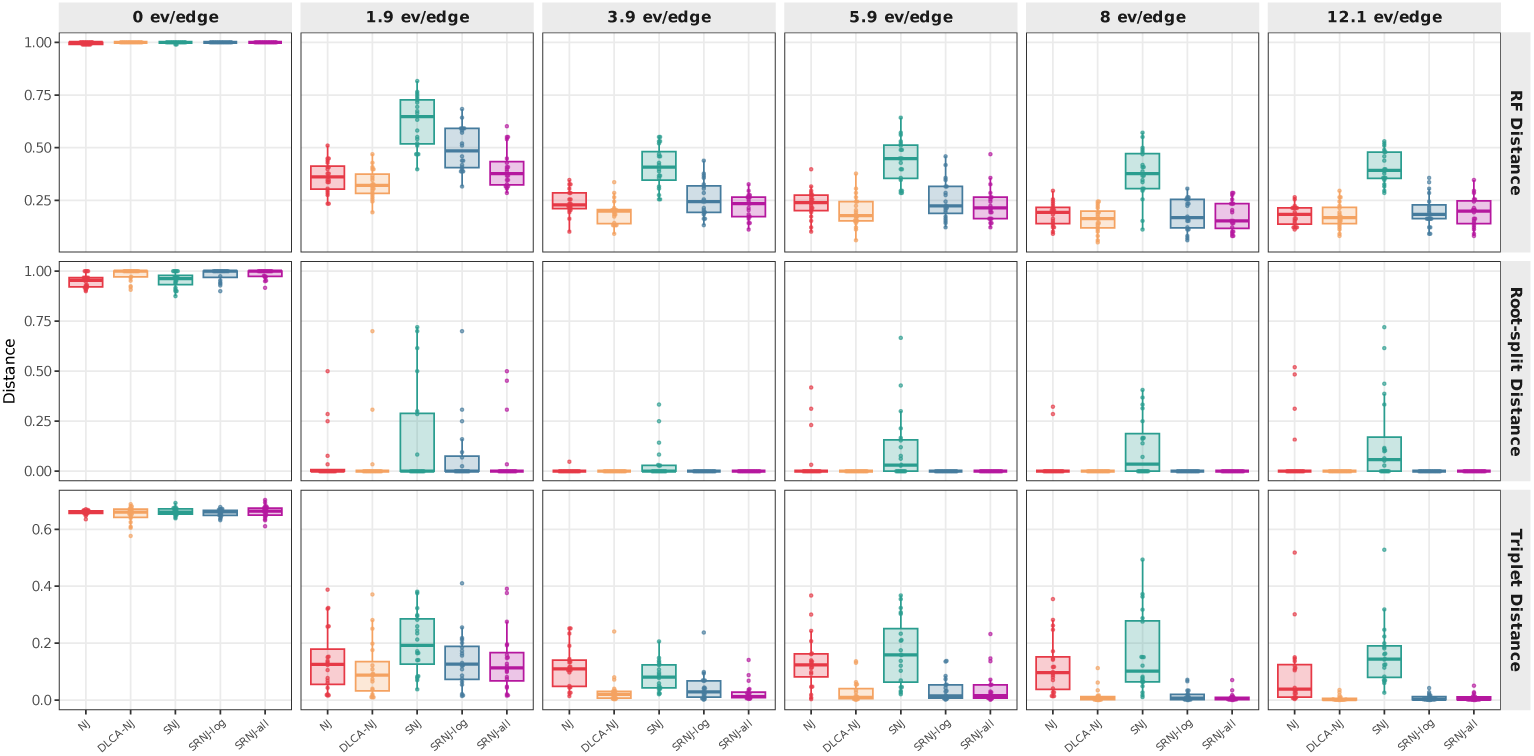
Effect of shared ancestral events on reconstruction accuracy. Reconstruction accuracy on *in-house* simulated data (same segment model as Fig. 2) as a function of the realised mean number of shared copy-number events per internal edge (left: star-like trees with mostly private, terminal events; right: strongly clonal trees with early, shared bursts), for *N* = 100 cells over 20 replicates. Rows show RF, root-split and triplet distance; lower is better. As shared ancestral signal increases, the LCA-based methods (DLCA–NJ, SRNJ) increasingly outperform NJ and SNJ, most clearly in the rooting-sensitive triplet and root-split distances.

### A.5 Simulation of CNAsim data

The realistic distance-estimation experiments (Figs 4 and A2) use datasets generated with CNAsim [39], which simulates read counts from a tumor copy-number evolution model and therefore allows the copy numbers to be inferred (by SCONCE2 or MEDICC2/HMMcopy) rather than assumed known. Figure A4 shows two example datasets. Each dataset contains, in addition to the tumor cells, a set of normal diploid cells; the copy-number profiles combine focal and chromosome-arm-level events, together with a strong founder amplification shared by all tumor cells.

#### Ground-truth control

On the in-house data the distances are computed from the known copy-number profiles and are therefore exact, whereas on the CNAsim data they are estimated by SCONCE2 from noisy reads. To determine whether the smaller advantage of the root-aware methods on CNAsim data (Fig 4) originates from the CNAsim generative model or from copy-number-inference error, we re-evaluated the same CNAsim datasets using ground-truth distances computed from the true (simulated) ancestral profiles, exactly as for the in-house benchmark. With these exact distances DLCA– NJ and SRNJ reconstruct the CNAsim trees accurately (RF ≈ 0.07 at *N* = 100), far better than the pairwise NJ and SNJ, mirroring the in-house result; once SCONCE2-nferred distances are used instead their accuracy degrades sharply (RF ≈ 0.4–0.6) and all methods become comparable. The muted CNAsim+SCONCE2 result is therefore driven by copy-number-inference error rather than by the CNAsim trees, separating the contribution of the tree reconstruction algorithm from that of the copy-number caller.

**Fig. A4.**
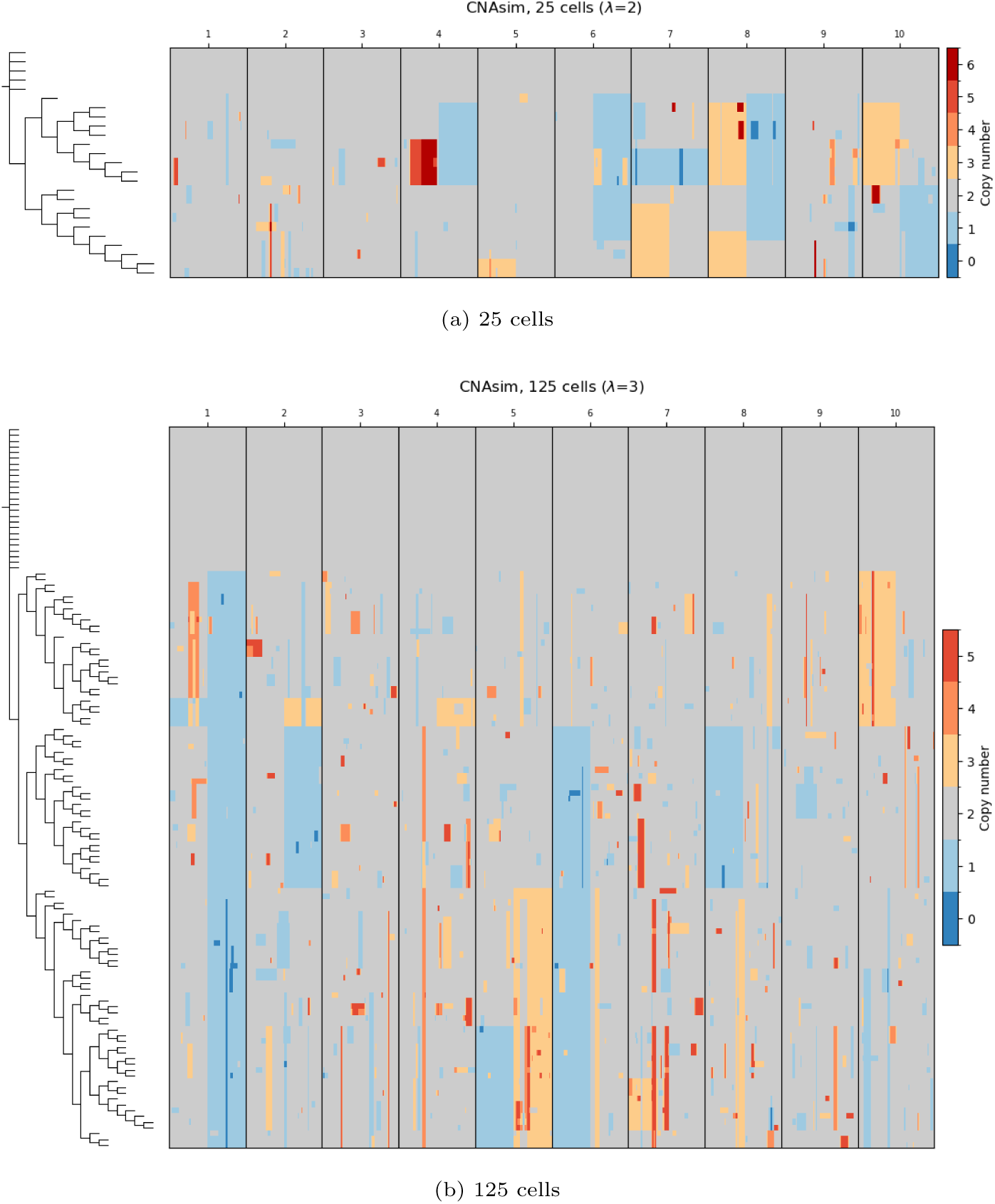
Example CNAsim-simulated tumor datasets. Example CNAsim-simulated tumor datasets with 25 (a) and 125 (b) cells, of which 20 and 100 respectively are tumor cells and the remainder are normal diploid cells (the ground-truth cell tree on the left, inferred copy numbers as a heatmap over the ten chromosomes).

